# Simultaneous Proportional Control of Two Degrees-of-Freedom Human Machine Interface Using Highly Sparse Sonomyography

**DOI:** 10.64898/2026.07.15.738689

**Authors:** Manikandan Shenbagam, Srikumar Venkataraman, Biswarup Mukherjee

**Author notes:** This manuscript was compiled on July 15, 2026. This work has been funded in part by the Science and Engineering Research Board, Department of Science and Technology, Government of India, through a Core Research Grant (CRG/2021/004967, PI: BM, Co-PI: SV), AO Foundation through a Asia Pacific National Research Grant (AOSRG2023009, PI:BM) and by the Department of Biotechnology, Government of India (BT/PR50638/MED/32/951/2023, PI: BM, Co-PI: SV).

## Abstract

Non-invasive human-machine interfaces (HMIs) are critical in developing prosthetic systems that offer intuitive, simultaneous, and proportional control over multiple degrees of freedom (DOFs). This study introduces a novel system for intuitive concurrent control of hand and wrist movements using sonomyography based imaging of muscle activity. Our method uses a sparse set of ultrasound scanlines to reduce computational complexity while enhancing usability. We evaluated four regression techniques for wrist and hand angle prediction, focusing on performance with a reduced sonomyographic feature set. We also explored the feasibility of a sonomyography-based system by simulating various factors that could affect prediction, including feature selection and scanline count. Our findings demonstrate that Gaussian process regression excels in predicting wrist and hand angles with just eight equispaced transducers in offline settings. Real-time evaluations with 10 non-disabled participants showed a 93 % success rate for two-DOF tasks using linear regression. The system was tested with an individual with amputation, achieving a 46 % success rate for two-DOF control in a 2D space, even though the ground truth data for model training was collected from the contralateral limb. This study validates our sonomyography-based approach for accurate wrist and hand angle estimation, reducing complexity and demonstrating potential in real-world scenarios.

## I. Introduction

**H**UMAN-Machine interfaces (HMIs) are pivotal for tracking upper limb positions and recognizing gestures, particularly in applications such as prosthetic control, remote manipulation, and virtual reality. It is widely acknowledged that these interfaces must be intuitive, robust, and portable, enabling natural movements and functioning effectively in various environments [1]–[3]. However, the rejection rate of upper limb prostheses remains high, approximately 44 %, mainly due to the lack of intuitive and advanced control systems, especially for managing multiple degrees of freedom (DoFs) simultaneously [4]. Most commercially available prosthetic hands offer a limited number of functions, controlled individually and not very intuitively [5], underscoring the need for more advanced and user-friendly control systems.

At present, surface electromyography (sEMG) is the primary technology used for HMIs to track upper limb positions and recognize gestures [6]–[10]. Nevertheless, sEMG-based HMIs face several limitations due to the stochastic and non-stationary nature of the sEMG signal, low spatial resolution, and susceptibility to crosstalk effects [11]. These limitations make it challenging to achieve proportional and simultaneous actuation over multiple DoFs. To overcome these challenges, ultrasound (US) and sonomyography have emerged as promising alternatives [12]. Unlike sEMG, ultrasound-based methods offer better mapping to hand kinematics by identifying muscle morphology independently of muscle activity. Sonomyography, which continuously monitors muscle activity using brightness mode (B-mode) ultrasound [13]–[15], has demonstrated higher performance compared to sEMG-based methods [16]. Moreover, sonomyography-based control interfaces provide better robustness making them more suitable for real-life applications [17], [18]. While previous studies have employed computationally intensive image processing operations for feature extraction from ultrasound B-mode sequences, recent advancements have made sonomyography more applicable for real-life HMI control applications. Sonomyography has been successfully utilized for prosthetic control and other biomechatronic devices, showcasing accurate classification of hand gestures and individual digit movements [19]– [23]. Additionally, sonomyography enables proportional control of prosthetic devices by continuously tracking upper-extremity joint angles and torques [24]–[26]. Recent studies have demonstrated that even with single-element transducers, sonomyography-based methods can accurately classify hand gestures and finger motions, making them highly promising for wearable HMI control interfaces [27]. Though ultrasound-based biomechatronic control interfaces have shown promise, most studies used commercial B-mode systems, limiting real-world use. To address this research gap [16], [28], [29] demonstrated that a sparse selection of scanlines from B-mode images can accurately classify hand gestures. This sparse representation effectively emulates the signals from a wearable ultrasound system by computing A-mode scanlines from the B-mode images. This technique has two distinct advantages: firstly, it enables testing and development of algorithms for wearable ultrasound systems without the need for wearable hardware. Second, it reduces the computational complexity associated with B-mode imaging-based data processing and prediction pipelines allowing real-time implementation with reduced latencies.

Furthermore, efforts are underway to explore different feature sets and modeling methodologies to enhance joint force and movement estimation using sonomyography [29]. The majority of these methods rely on high-dimensional feature sets and computationally complex models, which limit their real-time applicability and integration into practical, wearable systems. Using complex models could result in longer prediction times, impeding user’s ability to control the prosthesis without delay or intuitively [30], [31]. Additionally, most multi-degrees of freedom control modalities relies on sequential movements, whereas natural hand control involves simultaneous changes in degrees of freedom [32], [33]. For wearable and embedded prosthetic devices, it is crucial to prioritize models and features that are simple, computationally efficient, and hardware-friendly while retaining adequate control accuracy. Therefore, there is a pressing need to identify simple, low-dimensional features derived from ultrasound data which can be utilized to implement simultaneous control of multiple DoFs. Additionally, it is essential that the derived features utilize lightweight machine learning models that can deliver real-time performance on wearable hardware platforms. Furthermore, the data acquisition strategy itself plays a pivotal yet underexplored role in influencing model accuracy and system latency, which are critical for practical deployments.

In this paper, we propose a novel sonomyography-based method for continuous, online wrist angle and grasp completion estimation from a sparse set of ultrasound scanlines. We select a small subset of ultrasound scanlines from B-mode ultrasound image sequences obtained from the forearm of non-disabled individuals performing a target achievement task. This significantly reduces the dimensionality of the feature space and the computational cost involved in modeling the features. These scanlines also emulate A-mode signals derived from equivalent single-element transducers, making the system readily adaptable for portable, wearable sonomyography hardware. A key contribution of this work is the introduction and systematic evaluation of three distinct data collection paradigms — Horizontal Scan, Vertical Scan, and Centre-Out Scan — to investigate how acquisition strategy influences regression model performance for prosthetic control. This emphasis on acquisition strategy is particularly valuable for wearable system design, where sensor placement is constrained. Additionally, we compare the offline wrist angle prediction accuracy of four regression approaches — linear regression, support vector regression, ensemble bagged tree regression, and Gaussian process regression. By prioritizing simple models like linear regression, we demonstrate that competitive prediction accuracy can be achieved with significantly lower computational overhead, thereby facilitating practical, real-time implementation in embedded or wearable hardware. We systematically analyze the effect of the number of transducers, the data collection paradigm used, and the choice of features on wrist angle prediction accuracy. Following the offline experiments, we finalize the parameters and then validate the system online with non-disabled individuals as well as one individual with amputation.

## II. Materials AND METHODS

### A. Participants

We recruited 15 non-disabled participants for this study. 5 subjects (Mean age: 26 *±* 4 years) were initially recruited to perform offline pilot studies. Additional 10 participants (Mean age: 26 *±* 5 years) were recruited for the online validation of the proposed techniques. Written informed consent was obtained from all participants. Ethical approval for conducting the study was obtained from the respective institute ethics committee (IEC) at the Indian Institute of Technology, New Delhi (P021/P050). Individuals with diabetes, chronic pain, neuropathies, muscle contractures, and cognitive or neurological deficits were excluded from the study. We also recruited one individual with transradial amputation (Age: 19 years) at the All India Institute of Medical Sciences, New Delhi (AIIMS) after obtaining informed consent. The procedures were approved by the IEC at AIIMS (AHMSA00085/03.11.2023, RP-19/2023).

### B. Experimental procedures

Participants were comfortably seated with their dominant arm placed on a platform, aligning the elbow directly below the shoulder on a padded armrest. A high-frequency linear ultrasound probe (7.5 MHz, 0.3 mm pitch, 128 elements) was connected to a clinical ultrasound imaging system (ADC sampling rate 80 MHz, ADC resolution 12 bit, 32 channels, Surabi Biomedical Instrumentation Pvt. Ltd., India) and secured over the ventral aspect of the forearm muscle using a custom-made 3D printed ultrasound probe holder with velcro straps, as depicted in Fig. 1(a). Ultrasound gel (Sonocare transmission gel, Hi-tech Surgical & Fitness, India) was applied between the probe and the skin to facilitate imaging. The ultrasound probe was positioned transversely to image the major flexor muscles of the forearm, including the brachioradialis, flexor carpi radialis, flexor pollicis longus, and flexor digitorum superficialis, approximately 5 cm from the olecranon process. Participants wore a data glove sensor (5DT data glove, Fifth dimension technologies, USA) to measure the metacar-pophalageal joint angle of the digits, while a custom-designed inertial measurement unit (IMU) sensor (Model: Teensy LC board, Oregon, USA. with Adafruit 9-DOF Absolute Orientation IMU Fusion Breakout - BNO055 sensor) was attached to their palm to measure the wrist joint angle. The output from the data glove and IMU sensor served as ground truth data for model training. Image sequences from the ultrasound system were streamed to a custom-developed MATLAB interface (Version 2022a, Mathworks Inc., USA) on a laptop PC (Model: Intel Core i7 – 7700HQ, 32GB RAM, 4GB NVIDIA GeForce GTX 1040Ti) in real-time, with an approximate average frame rate of 20 frames per second. The MATLAB-based algorithm stored ultrasound images collected during the experiment, along with data glove sensor values and IMU sensor values, facilitating dataset preparation. Participants were unable to visually monitor their hands during the task, as this was a deliberate design choice intended to encourage reliance on proprioceptive feedback for cursor control.

**Fig. 1.**
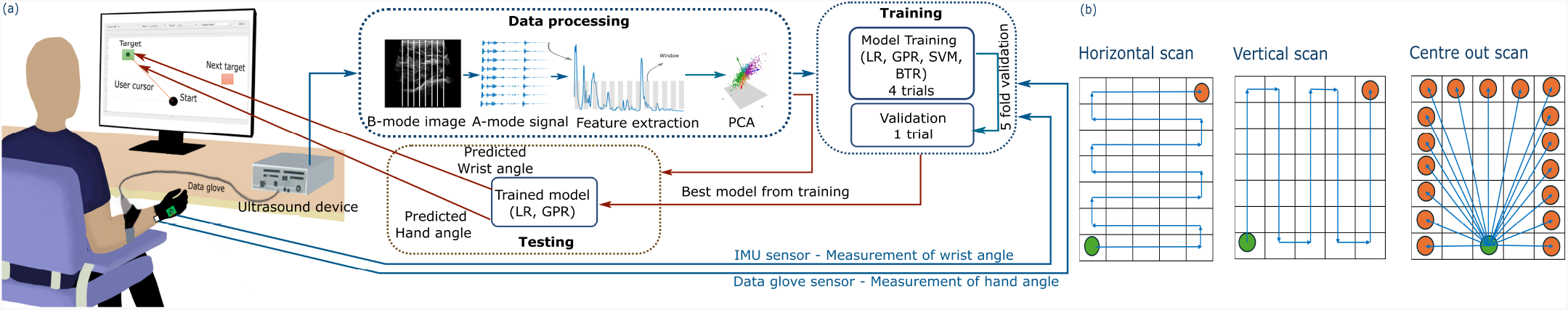
(a) Experimental setup showing volunteers equipped with an ultrasound probe on the forearm, a data glove, and an IMU sensor on the hand. The ultrasound system captures muscle activity and generates brightness-mode (B-mode) images. The data glove measures the hand’s opening angle, while the IMU records the wrist angle. Predicted values from the model are displayed as a black cursor on the screen. (b) Illustration of the data collection paradigms, including horizontal scan (HS), vertical scan (VS), and centre-out reaching scan (CO).

### C. Data processing

During the offline session, ultrasound images of the forearm, along with corresponding wrist joint angles and grasp completion measurements, were captured throughout the experiment and stored on the computer using MATLAB. During the calibration phase, the interface collected the maximum range of motion with the wrist supinated and pronated as well as the hand opened and closed. These maximum range values were further used to normalize the IMU and data glove measurements. Hand supination and pronation were mapped to the horizontal movement of the cursor on the interface, while hand open and close movements were mapped to the vertical movement of the cursor on the interface. During the rest position, when the hand was open and in a neutral position, the cursor was placed at the center bottom of the screen. Hand supination moved the cursor to the left side of the screen, and hand pronation moved it to the right side. Hand open placed the cursor at the bottom of the screen, and full hand close moved the cursor to the top of the screen. Targets are placed according to the data collection paradigm explained in Section II-D1. The aim of the user was to reach the target indicated in green. The next target to be achieved is then shown in green, guiding the user to move the cursor to reach it as shown in Fig. 1(b).

During this process, the MATLAB algorithm collected the normalized cursor position—horizontal movement from the IMU sensor output and vertical movement from the data glove output—along with the corresponding B-mode ultrasound images. A-mode captures one-dimensional echo amplitudes over depth, offering low computational cost and making it suitable for real-time, embedded applications. In contrast, B-mode generates two-dimensional grayscale images by mapping echo intensities across multiple scanlines, providing detailed anatomical information at a higher computational cost. So, the data processing pipeline, illustrated in Fig. 1, begins with the conversion of full-resolution B-mode images (775 × 607 pixels) from the forearm into sparse representations approximating A-mode signals. In the B-mode image, rows represent depth information and columns represent scanlines (channels). To reduce computational complexity, we evaluated performance using a reduced number of channels—specifically 4, 8, 12, and 16. For example, to reduce 607 channels to 8 channels, we calculated 607 / 8 = 75.8, rounding to 76 channels per group. However, dividing 607 columns into 8 groups does not result in equal-sized segments. Specifically, 607 / 8 = 75 with a remainder of 7. To address this, we distributed the columns as evenly as possible by assigning 76 columns to the first 7 groups and 75 columns to the final group. We then computed the mean across the columns within each group, resulting in 8 representative channels from the original 607 channels. We followed a similar procedure to reduce the channels to 4, 8, 12, and 16. This reduced form of the image is referred to as the sparse representation of the B-mode image. Therefore, the sparse image was reduced to 775 × 8. To compute the features from this data, we divided the depth information (i.e., 775 rows) into segments or depth-limited bins where each bin contained 50 points [29]. Specific features were extracted from each bin as discussed in Section II-D3. The calculated features from all channels of a single image are concatenated into a single vector, forming the feature matrix specific to that image. Following feature extraction, principal component analysis (PCA) was applied to reduce the dimensionality of the feature matrix in order to retain 95 % of the variance in the dataset. Post PCA, the feature matrix and corresponding ground truth values were fed into the machine learning model as shown in Fig. 1(a).

### D. Experiment 1: Offline model optimization

Our aim was to enable two degrees of freedom control with low computational cost and better accuracy. To achieve this, we optimized the regression models and tuned the parameters required to increase the model’s accuracy. In this section, we describe the protocol for model optimization and investigate the effect of several experimental and measurement factors on the performance of the regression models. For offline optimization of the model, data from five trials were collected in each data collection paradigm (Section II-D1). During the training phase, data from four trials were utilized, with the remaining trial reserved for model validation. A five-fold cross-validation procedure was employed for both training and validation. From this offline model training, the best machine learning model, optimal features, the required number of A-mode channels, and the most suitable data collection paradigm were determined. These parameters were subsequently utilized for the online training and testing protocol.

#### 1) Effect of data collection paradigm

We analyzed the effect of three different data collection paradigms on wrist and hand angle prediction accuracy as shown in Fig. 1(b). Participants were instructed that rotating their hand would control cursor movement in the horizontal direction, while opening and closing their fingers would control cursor movement in the vertical direction. In the horizontal scan, the users were guided by the interface to perform a raster scan where the horizontal direction was the fast axis and the vertical direction was the slow axis. The user started by rotating their wrist from rest to supination causing the cursor to move from the starting location at the bottom left to the box on the bottom right. Once the user reached the extreme points, they opened their hand by an appropriate amount to move to the next row, followed by continuing wrist rotation for horizontal traversal. Conversely, in the vertical scanning paradigm, participants initially move vertically, then rotate their hand to transition to the next column, and continue in the vertical direction. Both training paradigms enabled users to sequentially activate two degrees of movement: initially traveling in one degree before reaching the extreme position, then activating the second degree of movement. In the third case of center-out reaching movement, where targets are diagonally positioned from the origin, users were guided to simultaneously activate wrist and hand movements to travel in a straight line connecting the starting position and the intended target in order to reach the target. Guiding lines were shown to the user during the data collection process to help in traversing the path. For each target, the user travelled from the origin at the center (marked in green) to the respective target and then followed the same trajectory back to the origin.

#### 2) Effect of number of virtual transducers

We considered each ultrasound frame as a compilation of vertical scanlines. Each scanline can be thought of as an ultrasound transducer placed on the forearm and the resultant signal is an A-mode signal from the transducer, as shown in Fig. 1. Each scanline depicts a depth-resolved intensity map of the underlying anatomy, where peaks signify highly echogenic features such as bones and myofascial interfaces. To assess the impact of the number of transducers, we varied the number of scanlines from 4 to 16 in increments of 4. For each scanline, ultrasound features were calculated and employed in training and testing the regression models to examine the effect of the number of transducers.

#### 3) Effect of ultrasound image features

For each extracted scanline we applied a window of width 50 data points [29] along the depth of the scanline. Within each window, we extracted the following features: 1) Window mean, which calculates the average value within the window and represents the total acoustic energy reflected by the anatomical feature within it. 2) Window standard deviation, which measures the variability of values within the window and is thought to indicate the sharpness of the peak and, consequently, the thickness of the anatomical structure. 3) Window linear fit coefficients, obtained by fitting a straight line to the points within the window and calculating its slope and intercept. These coefficients also reflect the sharpness of the peak and the thickness of the anatomical structure. Additionally, we computed histogram features by constructing a histogram within the window with a bin width of 10. We analyzed the impact of these features extracted from the A-mode signal on the prediction accuracy. We trained all models with these features separately and reported the validation accuracy. Also we evaluated different feature combination strategies to assess their effect on model prediction accuracy; however, combining multiple features did not improve performance over mean features alone. Considering our goal of maintaining a low-complexity, real-time data processing pipeline, we prioritized mean features to minimize computational overhead and system latency.

#### 4) Effect of machine learning model

In our study, we utilized ultrasound features to evaluate the performance of four distinct supervised regression methods: simple linear regression (LR), ensemble bagged tree (BT), support vector regression (SVR), and Gaussian process regression (GPR). Through the application of these regression methods to our ultrasound features, we seek to determine the most effective approach for accurately predicting wrist and hand angles, considering each method’s unique characteristics and assumptions for our specific task

### E. Experiment 2: Online testing and validation with optimized models

During the online session, features from the ultrasound scanlines were calculated similarly to the offline session and tested with the best- and worst-performing models from the offline session: Gaussian process regressor and linear regressor. The ultrasound pipeline previously described was implemented online with a higher frame rate of approximately 20 frames per second and used to perform a two degrees of freedom simultaneous control test. As shown in Fig.,1, the blue cursor was controlled over the x-axis via the participant’s wrist pronation and supination (left to right) and over the y-axis via opening and closing of the hand. Participants were guided by visual cues and the experimenter to follow predefined training movements, spanning the 2 DOFs using the centre-out scan method. After training, participants were asked to reach 24 different targets spanning the entire 2D space as shown in supplementary Figure S2. The positions of these 24 targets were fixed, but the order in which the targets were presented to participants was fully randomized. Between targets, participants had to return to the center neutral position. To successfully complete a target, the participant had to stay within a normalized *±*5,% range of the target for a dwelling time of 1,s. If the target was not reached within 10,s, the trial was considered unsuccessful. The study followed a repeated measures design, such that all participants performed control tests with both evaluation models, allowing within-subject comparisons across methods.

### F. Evaluation metrics

Our primary outcome metric for assessing the performance of each regression model was the coefficient of determination (*R*^2^). *R*^2^ values were calculated to assess the prediction accuracy of both the predicted angle and the true angle in both hand and wrist angles.

We also computed the task-based outcome metrics to evaluate the performance of the user and the system in Experiment

2. We computed the success rate (in %) as the proportion of targets successfully acquired by the user:

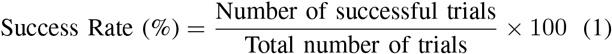

The movement time (in s) for each target was computed as the average time taken by the user to acquire the target from the starting or neutral point. We further computed the speed (in %/s) as the ratio of the path length (in %) to the movement time (in s) for each successful trial:

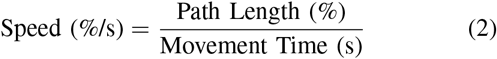

To assess movement quality, we evaluated the path efficiency (in %) traversed by the user in acquiring each target as:

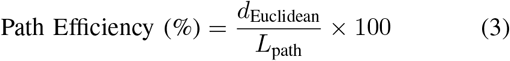

where *d*_Euclidean_ is the Euclidean distance between the start and end points, and *L*_path_ is the total length of the path, calculated by summing the Euclidean distances between consecutive points. Higher path efficiency indicates closeness to an idealized straight-line path connecting the target to the starting point.

Movement time, speed, and path efficiency were calculated only for successful trials. Failed trials were excluded from these outcome metric calculations to ensure that the results accurately reflect model performance during correctly completed tasks.

### G. Statistical tests

Due to the small sample size and the difficulty in verifying normality assumptions with multi-factorial parametric tests, we performed Friedman’s test to compare the performance of the regression methods. Pairwise comparisons, adjusted using a Bonferroni correction for multiple comparisons, were conducted to assess differences within factors affecting prediction accuracy. For the data collection paradigm and regressor model selection, where the factors were interdependent, we used two-way ANOVA to evaluate performance. We conducted a paired-sample t-test to compare the performance of the linear regressor and Gaussian process regressor for online experiments. All statistical analyses were performed using IBM SPSS Statistics (Version 28.0, IBM Corp, USA).

## III. Results

### A. Experiment 1: Pilot model training and parameter optimization

We conducted offline experiments to optimize various parameters including the machine learning model, data collection paradigm, number of channels, and type of features. Five non-disabled individuals were recruited separately for this purpose. The results of the parameter comparison are explained below. To assess the performance of different machine learning models, we fixed the number of channels at 8 and employed mean feature extraction across the length of A-mode signals, with bins of 50 in length. We trained a linear regressor, a Gaussian process regressor, a support vector machine, and a bagged tree regressor using four different data collection paradigms. The results indicate that the center-out scan (CO) and Horizontal Scan + Vertical Scan (HS+VS) paradigms outperformed the Horizontal Scan (HS) and Vertical Scan (VS) paradigms. The *R*^2^ values for different machine learning models across various data collection paradigms are illustrated in Fig. 2(a). The linear regressor achieved an *R*^2^ value of 0.76 *±* 0.18, the SVM achieved 0.74 *±* 0.09, and the bagged tree regressor provided 0.76 *±* 0.10. Notably, the GPR demonstrated superior performance with *R*^2^ values of 0.81 *±* 0.10. There were no statistical differences in the two-way interaction effect between data collection paradigms and machine learning models, nor were there differences between the machine learning models themselves.

**Fig. 2.**
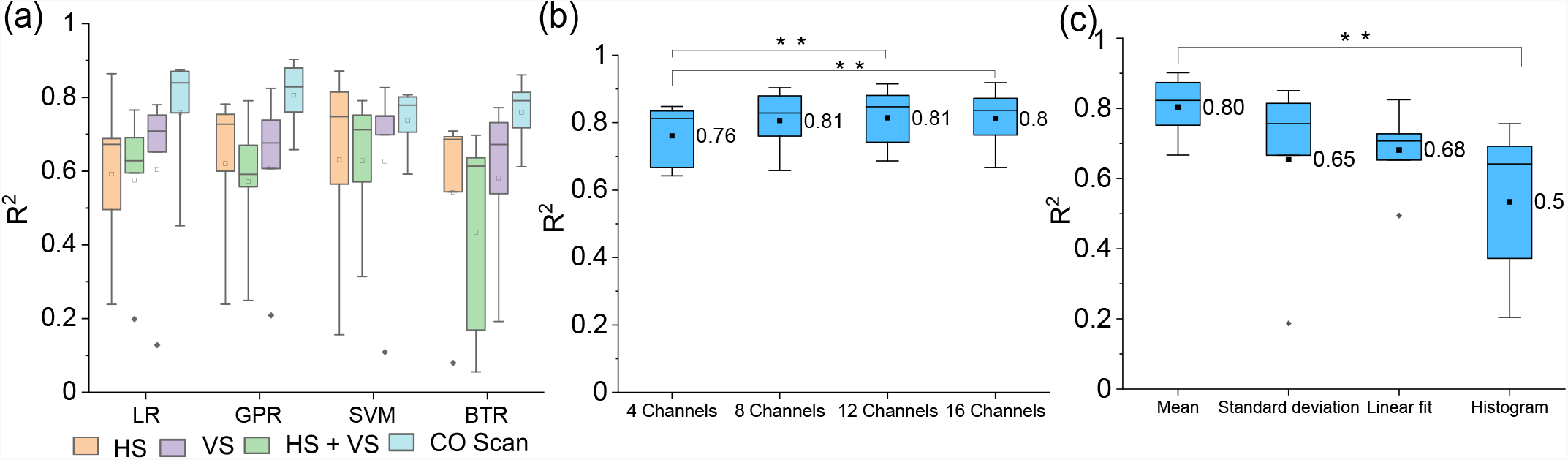
(a) Box plots showing the influence of data collection paradigm and machine learning model on the *R*^2^ coefficient for hand and wrist angle prediction. All models were trained with mean features extracted from eight equispaced scanlines. (LR — linear regression, GPR — Gaussian process regression, SVM — support vector regression, BTR — ensemble bagged tree regression, HS — horizontal, VS — vertical, HS+VS — horizontal and vertical combined, CO — centre-out scan). (b) Box plots depicting the influence of different numbers of channels on angle prediction accuracy. GPR models were trained with mean features extracted from 4, 8, 12, and 16 equispaced scanlines. (c) Box plots depicting the influence of different features extracted from each ultrasound scanline on angle prediction accuracy. GPR models were trained using mean, standard deviation, linear fit coefficients, and histogram features extracted from eight equispaced scanlines. Statistical comparisons were performed using Friedman’s test. Significance levels are indicated as follows: *p <* 0.05 (*), *p <* 0.01 (**), and *p <* 0.001 (***).

We also measured the data collection time for each paradigm. The data collection time for the horizontal scan was 137.81 *±* 40.05 s, for the vertical scan it was 160.02 *±* 70.77 s, and for the centre-out scan it was 316.82 *±* 134.46 s. The data collection time varied significantly across different paradigms, *χ*^2^(2) = 60.408, *p <* 0.001. Pairwise comparisons with a Bonferroni correction revealed significant differences between HS and CO Scan, HS and HS+VS, VS and CO Scan, and VS and HS+VS, with all *p*-values being less than 0.0001.

#### 1) Effect of number of channels

Given the superior results of GPR and CO scan in the previous section, we utilized the GPR model with CO scan data collection paradigm and mean features to evaluate the impact of different numbers of channels. The mean *R*^2^ values obtained were 0.76 *±* 0.10 for a 4 channels, 0.81 *±* 0.10 for 8 channels, 0.81 *±* 0.10 for 12 channels, and 0.81 *±* 0.1 for a 16 channels. As shown in Fig. 2(b), performance degradation was not observed beyond the 8-channel configuration. Prediction accuracy, was statistically different across the different numbers of channels selected for data collection, (p*<*0.005, *χ*^2^(8) = 10.680). Pairwise comparisons, performed with a Bonferroni correction for multiple comparisons, revealed that *R*^2^ values were statistically significantly different between the 4-channel and 16-channel systems (p = 0.042) as well as between the 4-channel and 12-channel systems (p = 0.020). However, the difference in prediction accuracy was not significant between 8-channel, 12-channel and 16-channel configuration (P=1.0). Therefore, there was no significant gains in prediction accuracy when the number of channels were increased beyond 8.

#### 2) Effect of feature selection on prediction performance

Based on the results from the previous tests, the GPR model with the CO scan as the data collection paradigm, and 8 channels, were used to investigate the influence of various features on model performance. We trained the GPR model using mean, standard deviation, linear fit coefficients, and histogram features derived from the scanlines of B-mode images. The corresponding *R*^2^ values are presented below. Specifically, the *R*^2^ values for the GPR model trained with mean, standard deviation, linear fit coefficients, and histogram features were 0.80 *±* 0.10, 0.65 *±* 0.27, 0.68 *±* 0.12, and 0.53 *±* 0.23, respectively, as shown in Fig. 2(c). Prediction accuracy, as measured by *R*^2^ values, was statistically significantly different across the different features selected to train the models, *χ*^2^(2) = 17.280, *p* = 0.02. Pairwise comparisons, performed with a Bonferroni correction for multiple comparisons, revealed that *R*^2^ values were statistically significantly different between histogram and mean features (*p* = 0.027).

### B. Experiment 2: Online validation and testing of the optimized models

Based on offline experiments, we optimized the data collection paradigm by selecting the CO scan, as it provided better accuracy compared to other paradigms. We observed that there was no performance degradation after 8 channels, so we fixed the number of channels at 8. Among the five features tested, the mean feature yielded the best accuracy. Therefore, for online experiments, these parameters were fixed. Our aim is to compare the performance of the linear regressor (second best performing but simplest) and Gaussian process regressor (best performing but complex) using these fixed parameters during online control. We hypothesized that with the humanin-loop, linear regression model performance will be at par with the Gaussian process regressor model.

Ten non-disabled individuals were recruited for online validation. Supplementary movies V1 and V2 demonstrate the process of user training and testing phases for non-disabled individuals. During the testing phase, targets appeared on the computer screen, and the user was prompted to attain the target using a cursor position predicted by the LR and GPR models, computed using the user’s dynamic muscle activity. We collected three trials of data for each participant using LR and three additional trials using GPR. A sample of the mean trajectory predicted by LR and GPR for one participant and selected targets is shown inFig. 3. The trajectories qualitatively indicate simultaneous activation of both DOFs for all the indicated targets across the entire 2D space. A quantitative assessment of the task performance based on the task-based outcome metrics have been presented next.

**Fig. 3.**
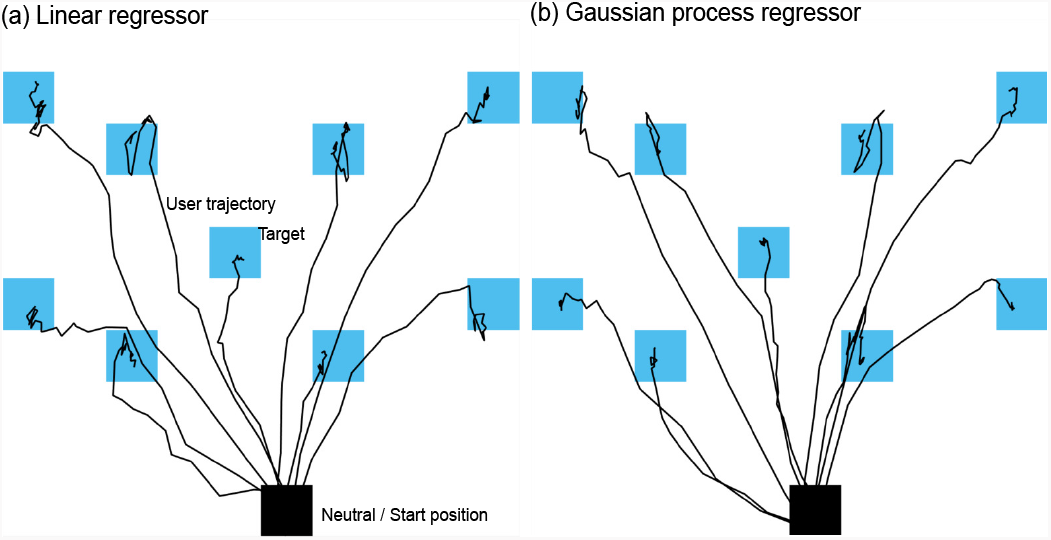
(a) Sample mean trajectory of 3 trials collected from an non-disabled volunteer and predicted by the linear regressor during online validation. (b) Mean trajectory of 3 trials collected from an non-disabled volunteer and predicted by the Gaussian process regressor during online validation.

#### 1) LR enables higher success rate

The success rate for individual targets across subjects is depicted in Fig. 4(a). The average success rate for the linear regressor was 93.08 *±* 8.18%, while for the Gaussian process regressor, it was 87.21 *±* 19.17%, as shown in Fig. 4(a). The success rate was statistically significantly different between the linear regressor and the Gaussian process regressor, *t*(23) = 2.23, *p* = 0.03, *d* = 0.45.

**Fig. 4.**
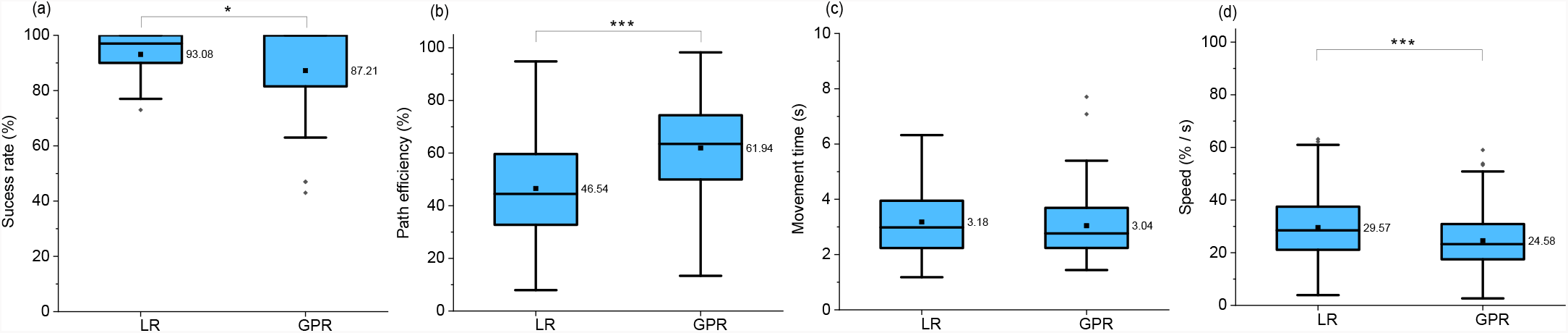
Box plots comparing the performance of the linear regressor and Gaussian process regressor across various outcome metrics, including (a) success rate, (b) path efficiency, (c) movement time, and (d) speed. Statistical comparisons were performed using paired t-tests. Significance levels are indicated as follows: *p <* 0.05 (*), *p <* 0.01 (**), and *p <* 0.001 (***).

#### 2) Higher path efficiency achieved with GPR

The path efficiency for individual targets is shown in Fig. 4(b) for both the linear regressor and the Gaussian process regressor. The average path efficiency for the linear regressor was 46.54 *±* 18.68 %, whereas for the Gaussian process regressor, it was 61.94 *±* 16.92 %, as depicted in Fig. 4(b). Path efficiency was statistically significantly different between the two regressors, *t*(597) = 18.07, *p <* 0.001, *d* = 0.73.

#### 3) LR enables higher movement speed but movement time is unaffected by the model

Fig. 4(c) shows the movement time, and Fig. 4(d) depicts the speed for both models across all targets. The average movement time for the linear regressor and the Gaussian process regressor was 3.18 *±* 1.25 s and 3.04 *±* 1.18 s, respectively, as shown in Fig. 4(c). The average speed for the linear regressor was 29.57 ± 11.49 %/s, and for the gaussian process regressor, it was 24.58 *±* 9.74 %/s, as shown in Fig. 4(d). The movement time was not statistically significantly different between the two regressors, *t*(70) = 0.67, *p* = 0.50, *d* = 0.08, while the speed was statistically significantly different, *t*(627) = 8.72, *p <* 0.001, *d* = 0.34

#### 4) Model training and prediction Time

The training time required for each model, as well as the prediction time for hand and wrist angles, is summarized below. The training times for the linear regressor and the Gaussian process regressor were 1.16 *±* 0.66 seconds and 45.28 *±* 9.13 seconds, respectively. Training time was statistically significantly different between the linear regressor and the Gaussian process regressor, *t*(9) = 15.95, *p <* 0.001, *d* = 5.04. The prediction times for the linear regressor and Gaussian process regressor were 0.57 *±* 0.21 ms and 4.80 *±* 2.35 ms, respectively. The prediction time was statistically significantly different between the two regressors, *t*(49959) = 401.4, *p <* 0.001, *d* = 1.8. usepackagegraphicx

#### 5) Pilot online validation with an individual with transradial amputation

The developed system was tested with an individual with transradial amputation. The participant had unilateral amputation and therefore, the contralateral intact limb was to obtain ground truth data for the hand and wrist angles. However, the ultrasound-imaging system was fixed to the amputated limb on the residuum using procedures similar to non-disabled participants as shown in Figure S1. The participant was instructed to mirror the intact limb’s motion with their imagined (phantom) right hand. The outcomes from the online trajectory, including success rate, path efficiency, movement time, and speed values, are presented in Supplementary Figure S3. Additionally, a 2D target achievement task performed by the amputee is shown in the supplementary video V3. The average success rate was 46 % for Linear Regression (LR) and 56% for Gaussian process Regression (GPR). Path efficiency was 37 % for LR and 41 % for GPR. The average speed was 30%/s for LR and 32%/s for GPR. The average movement time was 5.95 s for LR and 5.22 s for GPR. To further investigate the effects of using mirrored limb data as ground truth, we recruited 5 able-bodied subjects who performed the same task with ground truth data collected from their contralateral limbs. The ultrasound system was fixed similarly as in the case of the individual with amputation, and subjects were instructed to mirror the contralateral limb’s motion. The average success rates for these able-bodied subjects were 41.06 % *±* 17.98 % for LR and 46.4 % *±* 21.89 % for GPR—values comparable to those observed in the individual with amputation. However, from Section III.B.1, when able-bodied subjects performed the task with direct data collection (i.e., without mirroring), success rates were much higher: 93.08 % *±* 8.18 % for LR and 87.21 % *±* 19.17 % for GPR. These results suggest that errors introduced by the mirrored action during ground truth data collection can significantly degrade model performance. The success rate for each target is presented in Figure S4.

## IV. Discussions

In this study, we demonstrated real-time simultaneous and proportional control over two degrees of freedom using sparse ultrasound images obtained from the forearm flexor muscles. We showed that a sparse set of ultrasound scan lines obtained from B-mode ultrasound images enables the precise prediction of hand and wrist angles, thereby reducing the computational complexity of our model. Our technique compares the performance of the model using mean, standard deviation, linear fit coefficient, and histogram features extracted from sparse ultrasound images. Additionally, this study compares three different data collection paradigms with regard to regressor performance. The translational potential of this approach has been investigated in real-time experiments with non-disabled participants and with an individual with amputation. Also our proposed ultrasound-based system achieves a 93% success rate with an average movement time of 3.18 seconds, demonstrating comparable performance to state-of-the-art myoelectric control methods shown in Table I. Notably, while existing approaches like surface EMG, intramuscular EMG, and highdensity EMG achieve success rates between 85–99% with movement times ranging from 1.40 to 9.19 seconds, our system offers a noninvasive alternative with competitive realtime performance.

**TABLE I.** Comparison WITH State-OF-THE-Art Methods.

| Author | Dwell Time | Time to Achieve Target | Training Time | Target Threshold | Success Rate (%) | Sensing Modality | Machine Learning Used |
| --- | --- | --- | --- | --- | --- | --- | --- |
| Sgambato et al. [2] | 500 ms | 30 s | 150 s × 3 sessions + 5 min familiarization | $\pm 10\%$ | 93.75 | 2 DoF, A-mode Ultrasound | Linear Regression |
| Nowak et al. [34] | 0.3 s | 20 s | 36 s + 10 min familiarization | $\pm 15\%$ | 84 | 2 DoF, EMG | Ridge Regression |
| Proposed Method | 1 s | 10 s | 5 min | $\pm 5\%$ | 93.08 | 2 DoF, Sparse Ultrasound | Linear Regression |

### A. Training paradigm has an effect on prediction performance

The crticial observation from our study was that the prediction performance (in offline settings) was dependent on the choice of model and perhaps more importantly, the data collection paradigm. GPR outperformed all other models in offline validation. In this study we observed that the horizontal and vertical paradigms activated single degrees of movement at a time, while the CO scan promoted the simultaneous activation of two degrees of freedom, making this paradigm more intuitive than the previous two data collection paradigms. Since we used both hand and wrist angles, the CO scan paradigm activated more muscles during training. This simultaneous activation enabled superior performance over HS, VS as well as combined HS, VS training paradigms. However, our study also noted that the CO scan training takes considerably longer time compared to HS and VS which may result in fatigue for the user.

### B. Simple mean feature and 8-channels are sufficient for intuitive control

The offline study also revealed that even for two degrees of freedom control, we only needed an 8-channel system to control the cursor, as there was no performance degradation beyond that. Our prior study involving single degree of freedom isometric force prediction using a sparse sonomyography approach also corroborates this finding [29].

Prior studies by several other groups have used multiple features extracted from A-mode ultrasonic signals for proportional joint angle and force prediction [2], [17], [21], [29]. A systematic evaluation revealed that simple mean features extracted from the scanlines outperformed other features such as linear fit coefficients, standard deviation and histogram. This finding has significant implications for the translational potential of our proposed method since, mean features can be easily extracted in computationally resource-constrained settings.

### C. Linear regressor provides higher success rate and speed but with lower path efficiency

The online study reveals that the LR yields a higher success rate; using LR, the user can reach all targets in two-dimensional space. However, analysing the path efficiency, GPR has a higher path efficiency than LR. This indicates that while users may have difficulty in reaching extreme targets with the GPR model, it provides precise and accurate movements, leading to higher path efficiency. For nearby targets, both the success rate and path efficiency of GPR are higher than those of LR (see Supplementary Figure S2 (a), (b) and (c), (d)). Thus, using GPR, targets can be accessed very precisely and accurately. However, GPR-based models are computationally complex, requiring longer training as well as prediction times.

Interestingly, LR-based model outperformed GPR in all task-based outcome metrics in online testing except movement time. This finding is contrary to our offline validation results suggesting that for human-in-loop tests with visual feedback, the sensorimotor control of the operator may be able to compensate for the inaccuracies in the prediction. However, this compensation could lead to increased cognitive burden due to enhanced visual attentional resource allocation during the task.

### D. Limitations, practical considerations and usability of sparse sonomyography

While our approach effectively reduces computational complexity by using fewer ultrasound scanlines, this sparsity can limit the accuracy and precision of control over multiple degrees of freedom. One practical consideration is the trade-off between the number of scanlines and the quality of movement prediction, which necessitates careful calibration to ensure reliable performance. The usability of the system is also influenced by the data collection paradigms employed, with some methods proving more intuitive but potentially requiring longer setup times. In practical use, transducer shift relative to the trained position, like electrode shift in sEMG systems, can degrade performance. Ultrasound also depends on consistent gel application for proper coupling, which may limit longterm usability. Additionally, our study restricted upper arm motion, but forearm muscle images can change with elbow or shoulder movement. The weight and inertia of a prosthesis or carrying loads during daily tasks may further affect image consistency. Another important limitation is that this study did not investigate the stability of sparse sonomyography signals across multiple sessions or over extended periods. The effects of doffing and donning the system and the duration over which a single training session remains valid were beyond the scope of this work and should be addressed in future studies. Despite these limitations, the study highlights that with an optimized configuration—such as the use of eight equispaced transducers and appropriate regression methodologies—sparse sonomyography can offer a balance of efficiency and accuracy, making it a viable option for practical, real-time applications in prosthetic control. Another consideration is the calibration time associated with the current data collection method, which requires approximately 5 minutes in the online experiments. In future work, we aim to reduce this by exploring incremental modeling approaches and optimizing target selection.

## V. Conclusion

This study demonstrates the potential of a sonomyography-based approach to facilitate accurate and intuitive control of prosthetic systems, specifically for simultaneous hand and wrist movements. By utilizing a sparse set of ultrasound scanlines, we were able to significantly reduce computational complexity while maintaining high performance. Our findings indicate that Gaussian process regression models outperform traditional linear regression in predicting wrist and hand angles, particularly in scenarios with limited data from a small number of transducers. The real-time evaluations with non-disabled individuals further validate the feasibility of this approach, achieving a high success rate in two-DOF tasks. Overall, the proposed methodology offers a promising solution for enhancing the usability and effectiveness of prosthetic systems, with significant implications for real-world applications.

